# Location and mapping of the human rostromedial tegmental nucleus and associated midbrain inhibitory nuclei regulating dopamine neurons

**DOI:** 10.64898/2026.01.12.699117

**Authors:** Anastasia Filimontseva, YuHong Fu, Glenda M Halliday

## Abstract

Animal experiments reveal distinct GABAergic cell clusters within the dopaminergic midbrain regions in the rostromedial tegmental nucleus (RMTg) and retrorubral fields (RRF) that have yet to be clearly defined in humans. These neurons send prominent inhibitory projections to dopaminergic neurons in the substantia nigra and ventral tegmental area that impact motor, reward and threat processing. We have identified GABAergic RMTg and RRF cell clusters in 6µm formalin-fixed paraffin-embedded transverse human midbrain sections from ten control cases obtained from the Sydney Brain Bank using immunohistochemistry for GABA and tyrosine hydroxylase. We determined the location and cell size of RMTg and RRF GABAergic neurons, further mapping these cell cluster in transverse 50µm thick cresyl violet stained serial midbrain sections (every 750µm) from previously published controls (Halliday et al. 1990a). GABAergic neurons were cytoarchitecturally distinct, with the largest GABAergic neurons in the RRF, followed by RMTg neurons which were larger than GABAergic neurons in the well-defined interpedunclular nucleus (Kruskal-Wallis test, p<0.0001). RMTg and RRF GABAergic neurons first appear in caudal transverse midbrain sections approximately 38mm above the obex. RMTg moves rostrally and medially from underneath the decussation of the superior cerebellar peduncle to just lateral to the interpeduncular nucleus. The RRF cluster also moves rostrally and medially to the parabrachial pigmented nucleus (PBP) just under the red nucleus. The GABAergic neurons in RMTg and RRF/PBP that modulate dopamine neuronal excitability have distinct morphologies in humans. Identifying these inhibitory neurons is key to evaluating their role in neurodegenerative diseases.

## Introduction

The majority of somatodendritic inputs to dopamine (DA) neurons in the substantia nigra pars compacta (SNpc) are GABAergic (~40-70% of neurons)(Bolam and Smith 1990; Henny et al. 2012). These inhibitory afferents arise from a number of sources extrinsic to the SNpc (e.g. from the periaqueductal gray, dorsal raphe, lateral habenula, and basal ganglia)(Omelchenko and Sesack 2010; Beier et al. 2015; Nieh et al. 2015; Hjelmstad et al. 2013; Lerner et al. 2015; Evans et al. 2020; Vounatsos and Gittis 2025a), but also from GABAergic neurons interspersed within the midbrain DA regions.

The main midbrain DA regions are the SNpc, retrorubral fields (RRF) and ventral tegmental area (VTA), with the VTA cytoarchitecturally subdivided into the more lateral parabrachial pigmented nucleus (PBP), medial paranigral (PN) and parainterfascicular (PIF) nuclei, and midline interfascicular (IF), central linear (CL) and rostral linear (RL) nuclei (Fu et al. 2012). Across these regions there are approximately three times more DA neurons compared with GABAergic neurons in rodents and nonhuman primates, with more regional variation in DA compared with GABAergic neurons (Nair-Roberts et al. 2008; Kelly et al. 2022). Within these DA regions, the RRF has the highest proportion of GABA to DA neurons compared to other midbrain DA clusters (Nair-Roberts et al. 2008; Vounatsos and Gittis 2025a; Kelly et al. 2022). In contrast only around 25% of SNpc neurons are GABAergic, which is the lowest proportion among the main DA regions, and the VTA has a moderate population of GABAergic neurons (Nair-Roberts et al. 2008; Kelly et al. 2022). In rodents, the highest densities of GABAergic neurons in the VTA are in the caudal PN and PIF subnuclei (Nair-Roberts et al. 2008), although the rostromedial tegmental nucleus (RMTg) has not been identified in humans but in other species it is a GABAergic nucleus in ‘the tail of the VTA’ (Barrot et al. 2012).

Many GABAergic neurons intermixed within these DA regions are local interneurons. In contrast, GABAergic neurons broadly innervating midbrain DA neurons concentrate only in the RMTg and RRF (Barrot et al. 2012; Smith et al. 2019; Vounatsos and Gittis 2025b). The RMTg was initially defined in rodents with retrograde injections into the SNpc and VTA which reliably identified this concentration of GABAergic neurons (Smith et al. 2019; Jhou et al. 2009a; Lammel et al. 2012). In addition to its direct inhibition of the motor nigrostriatal DA pathway (Bourdy et al. 2014), the RMTg in general encodes negative prediction errors and plays a critical role in behavioural inhibition in general, as well as punishment learning through mesolimbic DA pathways (Jhou 2021). As discussed above, of the midbrain DA regions the RRF has the highest concentration of GABAergic neurons which densely innervate the SNpc, although their function is less well defined (Vounatsos and Gittis 2025b). In gerbils, the RRF GABAergic pathway is involved in male sex behaviour (Simmons et al. 2011) with RRF projections in general involved in reward processing, motivation and threat evaluation (Moaddab and McDannald 2021; Vounatsos and Gittis 2025b).

Due to species differences in midbrain anatomy and neural identification using developmental markers (e.g. some transcription factors for RMTg neurons are only expressed in embryonic tissue and not in adult human brain (Allen Institute for Brain Science 2010)), the human RMTg has yet to be defined and GABAergic clusters in the human RRF have largely been ignored. The fact that there are multiple GABAergic populations in close proximity within these anatomical regions (Margolis et al. 2012) further complicates the identification of these important GABAergic neuronal nuclei in the human midbrain. However, the comparative identification of these inhibitory neurons is important to determine as these midbrain nuclei provide broad inhibition to midbrain DA neurons with any deficits likely to have significant impact.

For the first time we mapped the RMTg and a large RRF GABAergic neuronal cluster (LatC) in the human brainstem using immunohistochemistry and three-dimensional reconstructions in serial sections showing that these GABAergic cell clusters have distinct morphological features and anatomical locations. Overall, we present detailed anatomical maps of these prominent human midbrain GABAergic nuclei that innervate midbrain DA neurons.

## Materials and Methods

### Human brain tissue

For immunohistochemical analysis, formalin-fixed paraffin-embedded (FFPE) cross-sectional midbrain tissue blocks from aged healthy controls without neurological or neuropathological diseases were obtained from the Sydney Brain Bank (n=10, mean age[standard deviation]=85[12] years, for details see Supplementary Table 1). Serial section reconstruction using thick cross-sectional Nissl-stained midbrain sections from control cases (n=4, for details see Supplementary Table 1) previously processed and published (Halliday et al. 1990b) were also included. These midbrain sections were cut at 50μm on a freezing microtome, collected at 750μm intervals, and stained with 0.5% cresyl violet (HB5608 Hello Bio), as previously described (Halliday et al. 1990b). The study was approved by the University of Sydney Human Research Ethics Committee (2021/845).

### Immunohistochemistry

FFPE tissue blocks were sectioned at 6µm and dried on microscope slides at 37°C for 48 hours. Sections were deparaffinized by incubation at 60°C for 1 hour and then submerged in D-Limonene (Hurst Scientific, HISTO-5LCTN) for 2 x 7 minutes, followed by rehydration in decreasing ethanol concentrations (100% ethanol for 2 x 3 minutes, 95% ethanol for 3 minutes, 70% ethanol for 3 minutes) and distilled H_2_O for 3 minutes. Sections underwent heat-induced antigen retrieval (HIAR) in commercially available antigen retrieval solution (Invitrogen, 00-4956-58) diluted to 1x using a programmable antigen retrieval cooker (Aptum Biologics Ltd, UK, Retriever 2100). Peak temperature reached ~121°C and afterwards sections were gradually cooled for two hours. Sections were washed with tris-buffered saline (TBS, Sigma-Aldrich, 94158-10TAB), and endogenous alkaline phosphatase (AP) and hydrogen peroxidase (HRP) were quenched with BLOXALL blocking solution (SP-6000-100) for 30 minutes. Afterwards sections were blocked for 1 hour at room temperature with in-house made normal horse serum blocking solution (2% horse serum, 1% BSA in 0.01M TBS). Sections were incubated with a primary cocktail of antibodies to tyrosine hydroxylase (TH, rabbit polyclonal IgG, AB152 Sigma-Aldrich, RRID:AB_390204, diluted 1:200) and GAD67 (mouse monoclonal IgG_3_, sc-28376 Santa-Cruz Biotechnology, RRID:AB_627650, diluted 1:50) for 4 nights at 4°C. Secondary antibody incubation for 90 minutes at room temperature used a cocktail of ImmPRESS™-alkaline phosphatase (AP) horse anti-mouse IgG polymer detection kit (MP-5402 Vector Laboratories, RRID:AB_2336535) for GAD67 and ImmPRESS™-horseradish peroxidase (HRP) horse anti-rabbit IgG polymer detection kit (MP-7401 Vector Laboratories, RRID:AB_2336529) for TH. Antibody detection was visualised using SignalStain® Vibrant Red AP Substrate Kit (76713 Cell Signaling) or ImmPACT® DAB EqV HRP Substrate Kit (SK-4103 Vector Laboratories) prior to standard dehydration in increasing graded ethanol concentrations and clearing in xylene, then mounting with Eukitt® quick-hardening mounting medium (03989 Sigma-Aldrich). In separate control experiments where either the primary or secondary antibodies were omitted, no AP or HRP was detectable.

### Image acquisition

An automated slide scanner (SLIDEVIEW VS200, Tokyo, Japan) was used to scan the sections using pre-set focusing and exposure parameters. The method for brightfield image acquisition has previously been described (Filimontseva et al. 2025) with slight amendments to optimize the exposure settings for 1) the GABA and TH stained sections, and 2) for serial Nissl stained sections.

### Identification of GABAergic cell clusters

To identify RMTg, RMTg published anatomical maps from rodents and non-human primates were used (Supplementary Figures 1-3) and the most frequent landmarks determined (Supplementary Table 2). GABA and TH stained thin sections were used to identify GABAergic neuronal clusters within the dopaminergic regions based on their morphology, orientation, density and intensity of GABAergic staining (see Figure 1). GABAergic cell clusters were compared to already identified cell clusters in the adjacent interpeduncular nucleus. To ensure differentiation of neurons in these cell clusters, a semi-automated method was used to perform morphological analysis of the identified GABAergic neuronal clusters. For a full description see Supplmentary Methods. Briefly, images of Nissl stained and GABA stained sections were loaded into QuPath (ver. 0.5.1, RRID:SCR_018257, https://qupath.github.io/) (Bankhead et al. 2017) and the system was trained to identify neurons in the region of interest based on stain intensity. Successful training was used to create a pixel classifier and identified neurons were filtered by size and shape (Supplmentary Methods). Due to technical variations in the intensity of GABA staining on the human FFPE tissue, this analysis was restricted to six cases with similar conspicuous GABA neuronal staining. The morphological features of these GABAergic neurons were compared to Nissl stained neurons in the same regions analysed between 33mm to 38mm Obex at every 1.6mm interval.

**Figure 1.**
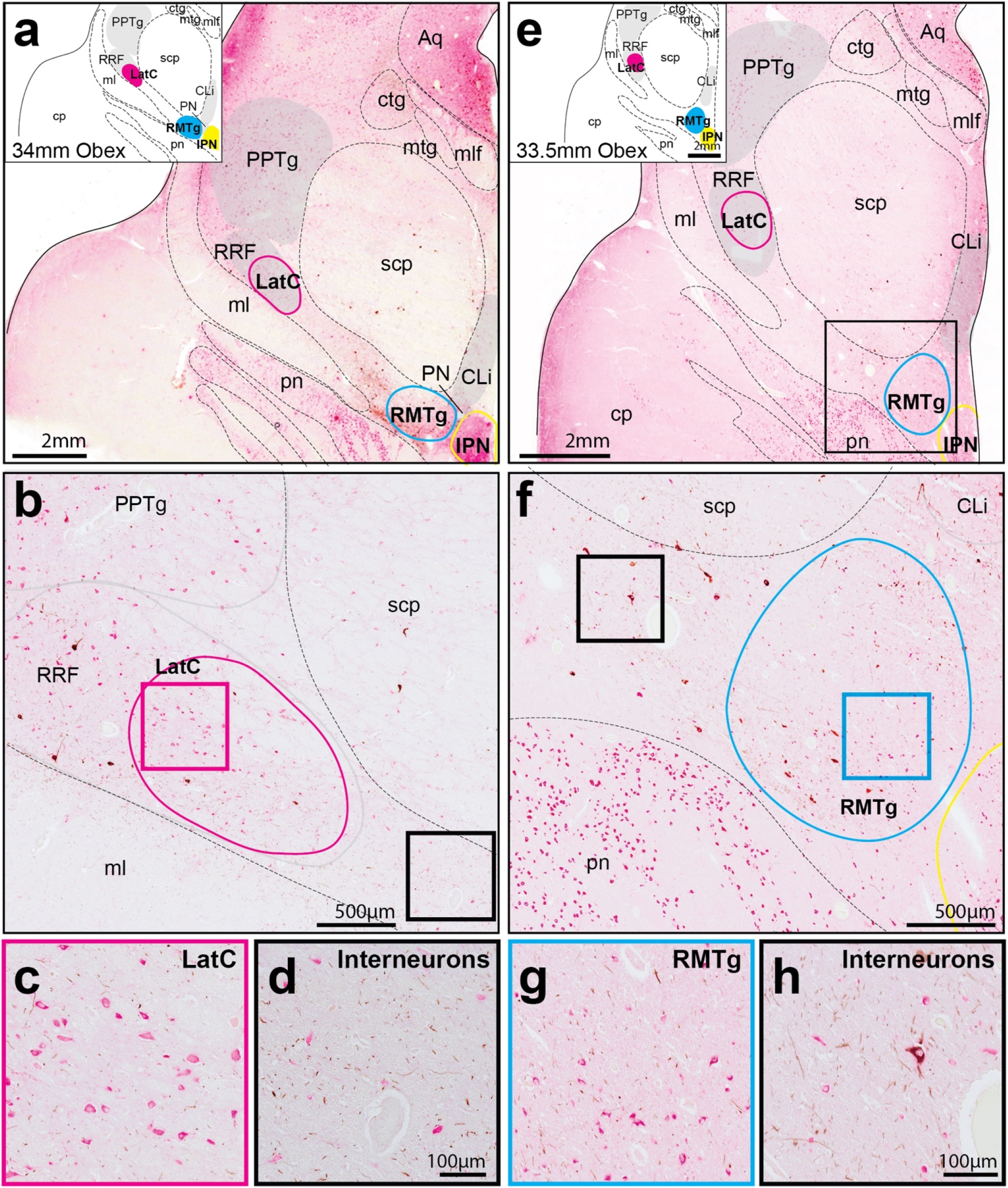
Detailed illustration of novel cluster identification in GABA (pink alkaline phosphatase) and TH (brown diaminobenzidine) immunohistochemically stained sections. Schematic chemoarchitectural maps were generated by tracing key landmarks and nuclei in the caudal midbrain (a, e). At these levels, there was a lateral cluster (LatC) embedded within the retrorubral field (RRF)(b) with the rostromedial tegmental nucleus (RMTg) located dorsolateral to the interpeduncular nucleus (IPN) and ventral to the superior cerebellar peduncle (scp)(e, f). GABAergic RMTg neurons were identified by their larger size and more intense staining from neighboring interneurons (g, h). Higher magnification images revealed distinct morphology, orientation and density that differentiates LatC neurons from neighboring regions (c). These LatC neurons also appeared different from fainter, more scattered inhibitory interneurons (d). Scale bars equal 2mm in a and e, 500μm in b and f, and 100μm in c, d, g, and h. Other abbreviations: aqueduct (Aq); caudal linear nucleus (CLi); cerebral peduncle (cp); central tegmental tract (ctg); mammillotegmental tract (mtg); medial lemniscus (ml); medial longitudinal fasciculus (mlf); paranigral nucleus (PN); pedunculopontine tegmental nucleus (PPTg); and pontine nuclei (pn).

### Anatomical mapping

Midbrain DA regions were cytoarchitecturally mapped on digitized images in Adobe Photoshop (ver. 25.9.1, Adobe Systems Inc., San Jose, CA, USA, RRID:SCR_014199, https://www.adobe.com/products/photoshop.html) and Illustrator (ver. 28.5, Adobe Systems Inc., San Jose, CA, USA, RRID:SCR_010279, http://www.adobe.com/products/illustrator.html) based on the human brainstem atlas and previously published cytoarchitecture studies (Paxinos et al. 2020; McRitchie et al. 1995; McRitchie et al. 1997). For the GABA and TH stained sections, TH staining was used to determine the DA brain regions and for the serial Nissl stained sections, neuromelanin pigment was used to determine the DA brain regions. GABAergic neuronal clusters were mapped in all cases using the GABA and TH stained sections and the regions identified in the serial Nissl sections. Both thick and thin sections were used to reconstruct proposed three-dimensional models of the DA and GABA regions of interest. The midline and aqueduct were used to align every other 50μm, taken at 750μm intervals. Landmarks were outlined and relevant clusters colour coded in Adobe Photoshop. Stacked Nissl and GABA stained sections were compiled in Adobe Illustrator, and mapped structures compared.

### Analysis of cell types within GABAergic cell clusters

For this analysis the small numbers of dopaminergic neurons that were observed on the borders of GABAergic clusters were ignored. The GABA and TH stained sections were decoverslipped, rehydrated and counterstained with 0.5% cresyl violet (HB5608 Hello Bio). Sections then underwent standard dehydration in increasing graded ethanol concentrations and cleared in xylene, prior to mounting with Eukitt® quick-hardening mounting medium (03989 Sigma-Aldrich). Within the mapped borders of the GABAergic cell clusters, the numbers of GABA and non-GABA Nissl positive neurons were counted and their densities and proportions calculated.

### Statistical analysis

Statistical analyses were performed in GraphPad Prism (version 10.0.3, RRID:SCR_002798 GraphPad Software Inc., California, United Stated). Kruskal-Wallis non-parametric test was used to compare cell circularity of neurons in the GABAergic cell cluster of the RMTg, RRF and interpeduncular nucleus in Nissl and GABA stained sections. A two-way ANOVA was run to analyse effect of anatomical region and tissue stain on neurnal size, followed by post-hoc Tukey tests. The proportion and density of GABA to Nissl neurons in the GABAergic cell clusters was also analysed with two-way ANOVA tests. For all analysis the statistical significance was set to p-value < 0.05.

## Results

### Comparative landmarks to identify the human RMTg

We identified comparative cytoarchitectural landmarks for the human RMTg based on animal mapping literature (Supplementary Table 2). These important landmarks from this literature oriented our anatomical search for the RMTg in human midbrain tissue sections (Supplementary Table 2). The RMTg in other species first appears at the level of caudal interpeduncular nucleus, posterior to PN (Supplementary Figure 1)(Jhou et al. 2009b; Kaufling et al. 2009). As the superior cerebellar peduncle appears, the RMTg shifts dorsally (Supplementary Figure 2). The RMTg is elongated at this level where it shifts laterally below the RRF.

### Morphological identification of distinct inhibitory cell clusters in the human midbrain DA regions

Assessment of human GABAergic cell clusters using the comparative landmarks from the literature revealed two clusters in the caudal midbrain, the RMTg located close to the interpeduncular nucleus and another cluster slightly more lateral (LatC)(Figure 1). To determine if these two clusters were distinct morphologically, we compared their GABAergic neurons using morphological analyses and also compared them to the GABAergic neurons in the interpeduncular nucleus. GABAergic neurons in these cell clusters appeared to be morphologically different (Figures 1 and 2a, b), with the RRF LatC GABAergic neurons larger, rounder and more densely labelled compared to the small-medium sized RMTg GABAergic neurons that stained lighter and were randomly oriented (Figures 1 and 2a, b). When measured (Figure 2c-f), the size of GABAergic neurons in these cell clusters were significantly different [F (2,10796) = 166.9, p<0.0001]. Post-hoc Tukey tests revealed that LatC GABAergic neurons were significantly larger than RMTg GABAergic neurons, with both being significantly larger than GABAergic neurons in the interpeduncular nucleus (Figure 2g). Analysis of neuronal size in Nissl versus GABA stained sections (Figure 2c-f) revealed a significant main effect of the stain used on neuronal size [F (1,10796) = 21.41, p<0.0001]. Further, there was a significant difference in neuronal size between anatomical region depending on the stain used [F (2,10796) = 31.60, p<0.0001], although GABAergic neurons in both the RMTg and LatC GABAergic neurons were larger on average than Nissl stained neurons (Figure 2g). Cell circularity measurements confirmed that LatC neurons were on average rounder than RMTg neurons [Kruskal-Wallis test, p<0.0001](Figure 2h), although not so much for the GABAergic neurons in these regions (Figure 2i). Overall, the size and morphology of these GABAergic neurons in both tissue preparations are reliable measures to identify these distinct midbrain GABAergic cell clusters.

**Figure 2.**
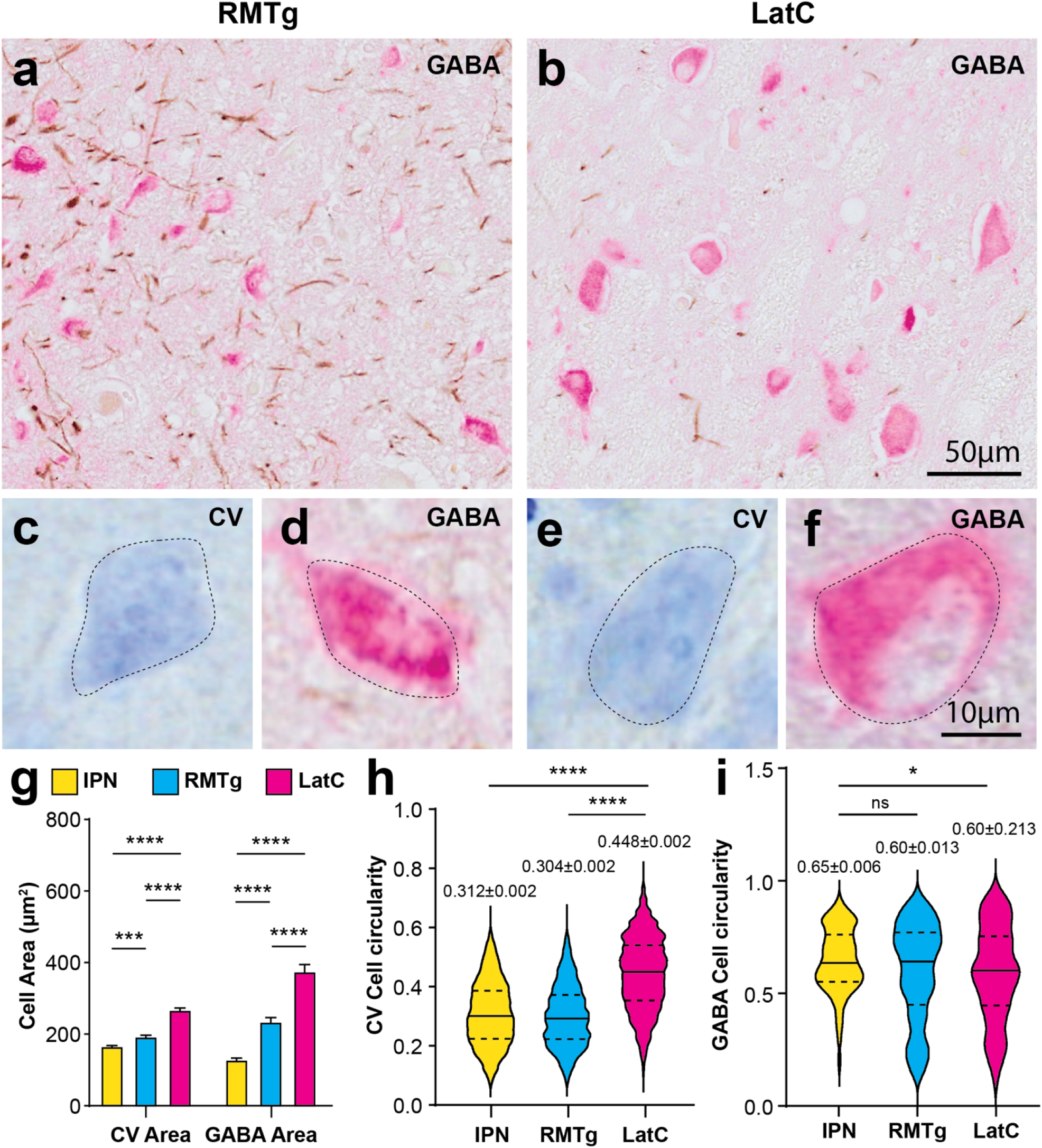
Morphological identification of distinct inhibitory clusters in the human midbrain dopaminergic regions. Micrographs are representative images from GABA and TH immunohistochemically stained sections (GABA labeling with alkaline phosphatase in pink; TH labeled with diaminobenzidine in brown) (a, b, d, e) showing specific neuronal clusters and high magnification of CV (c, e) and GABAergic neurons (d, f) with cellular annotations for morphology analysis overlayed. The rostromedial tegmental nucleus (RMTg) is presented in a, c, d. A novel, lateral cluster (LatC) of GABAergic neurons in the RRF is presented in b, e, f. Measures of neuronal area (g) and cell circularity (h, i) in CV (N=4) and GABA (N=6) stained sections validate the identification of these distinct inhibitory neuronal cell clusters in comparison to the interpeduncular nucleus (IPN, yellow = IPN, blue = RMTg, pink = LatC).. For area analysis, a two-way ANOVA revealed that anatomic region [F (2,10796) = 166.9, p<0.0001] and stain type significantly affected neuronal size [F (1,10796) = 21.41, p<0.0001] in g). Error bars display SEM. Post-hoc Tukey tests showed that LatC neurons were the largest, **** p<0.0001 and IPN neurons were the smallest [*** p<0.0001 in g]. GABAergic neurons in the RMTg and LatC were slightly larger than Nissl stained neurons. Cell circularity analyses also identified differences in these neuronal cell clusters in both tissue preparations. Violin plots revealed that LatC neurons were significantly rounder than RMTg and IPN in CV stained sections [Kruskal-Wallis test, **** p<0.0001 in (h)]. GABA staining revealed significant differences only between LatC and IPN circularity [Kruskal-Wallis test, * p<0.05 in (i)].

### Chemoarchitectural mapping of GABAergic cell clusters in midbrain DA regions

Midbrain DA regions surrounding the GABAergic RMTg and RRF LatC were cytoarchitecturally delineated and mapped using the GABA and TH colabelled sections (Figure 3). The RMTg first appeared caudally in the midbrain approximately 33-34mm above the Obex where it is at its largest in cross section mostly due to an absence of surrounding DA neurons and subregions (Figure 3a, b, 33.5mm, 34mm Obex). It lies lateral to the interpeduncular nucleus, dorsal to the pontine nuclei and ventromedial to the superior cerebellar peduncle (Figure 3a, b, 33.5mm, 34mm Obex). At these caudal levels, there is an obvious RRF LatC GABAergic cell cluster located some distance from the RMTg at the medial edge of the RRF on the ventrolateral edge of the superior cerebellar peduncle (Figure 3a, b, 33.5mm, 34mm Obex). In more rostral midbrain sections, the RMTg GABAergic cell cluster maintains its position but is reduced in size due to its location within the DA neurons of the VTA PN subregion and ascending DA fibres (Figure 3a, b, 34.5mm, 35mm, 35.5mm Obex). At 37mm Obex the RMTg appears as a small structure (Figure 3a, b, 37mm Obex) and discontinues in more rostral sections due to the appearance of the more dense compact SNpc clusters (beyond 37mm above Obex). The RRF LatC cell cluster maintains its position laterally (Figure 3a, c, 33.5mm, 34mm, 34.5mm, 35mm Obex) until the VTA PBP subcluster begins to expand laterally in more rostral sections (Figure 3a, c, 35.5mm, 37mm Obex). At these levels (35-36mm above the Obex) this GABAergic cell cluster bridges between the medial RRF and lateral PBP (Figure 3a, c, 35.5mm, 37mm Obex). With the discontinuation of the RRF rostrally, the LatC moves ventromedially to embed into the PBP on the ventral aspect of the superior cerebellar peduncle and continues to move ventromedially to be adjacent to the RMTg within the VTA DA regions (Figure 3a, c, 37mm Obex).

**Figure 3.**
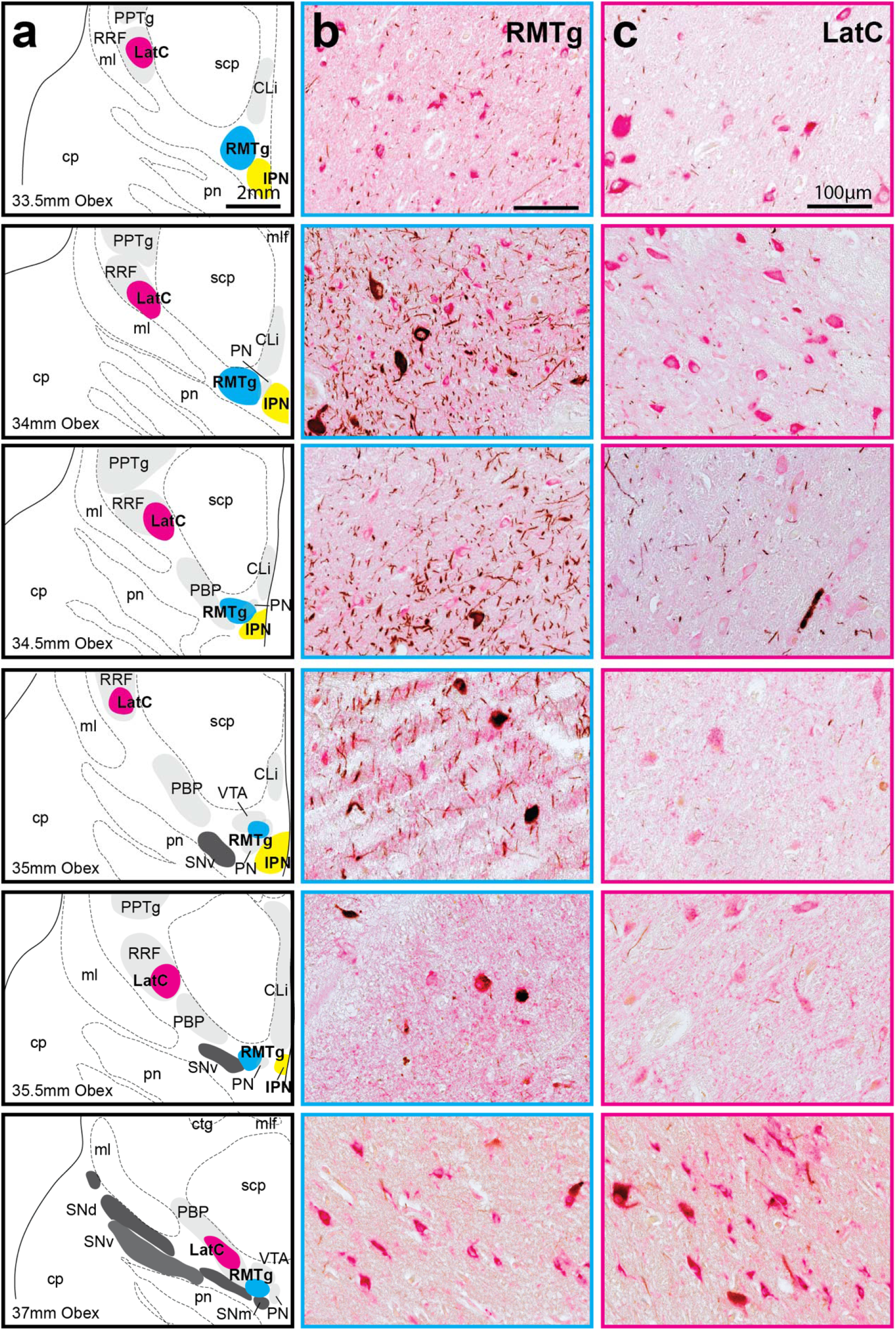
Chemoarchitectural mapping of GABAergic cell clusters in midbrain DA regions. Representative micrographs of GABA and TH immunohistochemically stained sections (GABA labeling with alkaline phosphatase in pink; TH labeled with diaminobenzidine in brown) show neurons in the RMTg in b and RRF/PBP LatC in c with maps of their location and the main DA regions shown in a. The levels mapped are identified relative to the obex. TH+ neurons were found in the dorsal, ventral and medial substantia nigra pars compacta (SNpc) tiers (SNd, SNv and SNM), retrorubral field (RRF), parabrachial pigmented nucleus (PBP), paranigral nucleus (PN), ventral tegmental area (VTA), caudal linear nucleus (CLi) and the interfascicular nucleus (IF). Other relevant landmarks include the central tegmental tract (ctg), cerebral peduncle (cp), interpeduncular nucleus (IPN), mammillotegmental tract (mtg), medial lemniscus (ml), medial longitudinal fasciculus (mlf), pedunculopontine tegmental nucleus (PPTg), pontine nuclei (pn) and superior cerebellar peduncle (scp). Main TH+ SNpc subclusters are annotated in dark grey, whereas VTA subclusters and other relevant landmarks are shaded in light grey. The nuclei of interest were mapped in the 6μm thick transverse midbrain sections along a 4mm distance in 10 cases. Scale bars equal 2mm in a and 100μm in b and c.

### Cytoarchitectural identification of the RMTg and RRF LatC in midbrain DA regions

The RMTg and RRF LatC were cytoarchitecturally identified and mapped using serial thick Nissl stained cross-sectional midbrain sections to observe cell clusters more readily through a greater number of midbrain levels (Figure 4). Midbrain DA regions were identified using neuromelanin pigment. The RRF/PBP LatC was observed within the same anatomical location in these midbrain DA subregions, that is, at the medial edge of the RRF (Figure 4a, b, c, 33mm, 34mm, 35mm, 36mm Obex) shifting in rostral sections to the dorsolateral PBP subregion (Figure 4a, b, c, 37mm, 38mm Obex).

**Figure 4.**
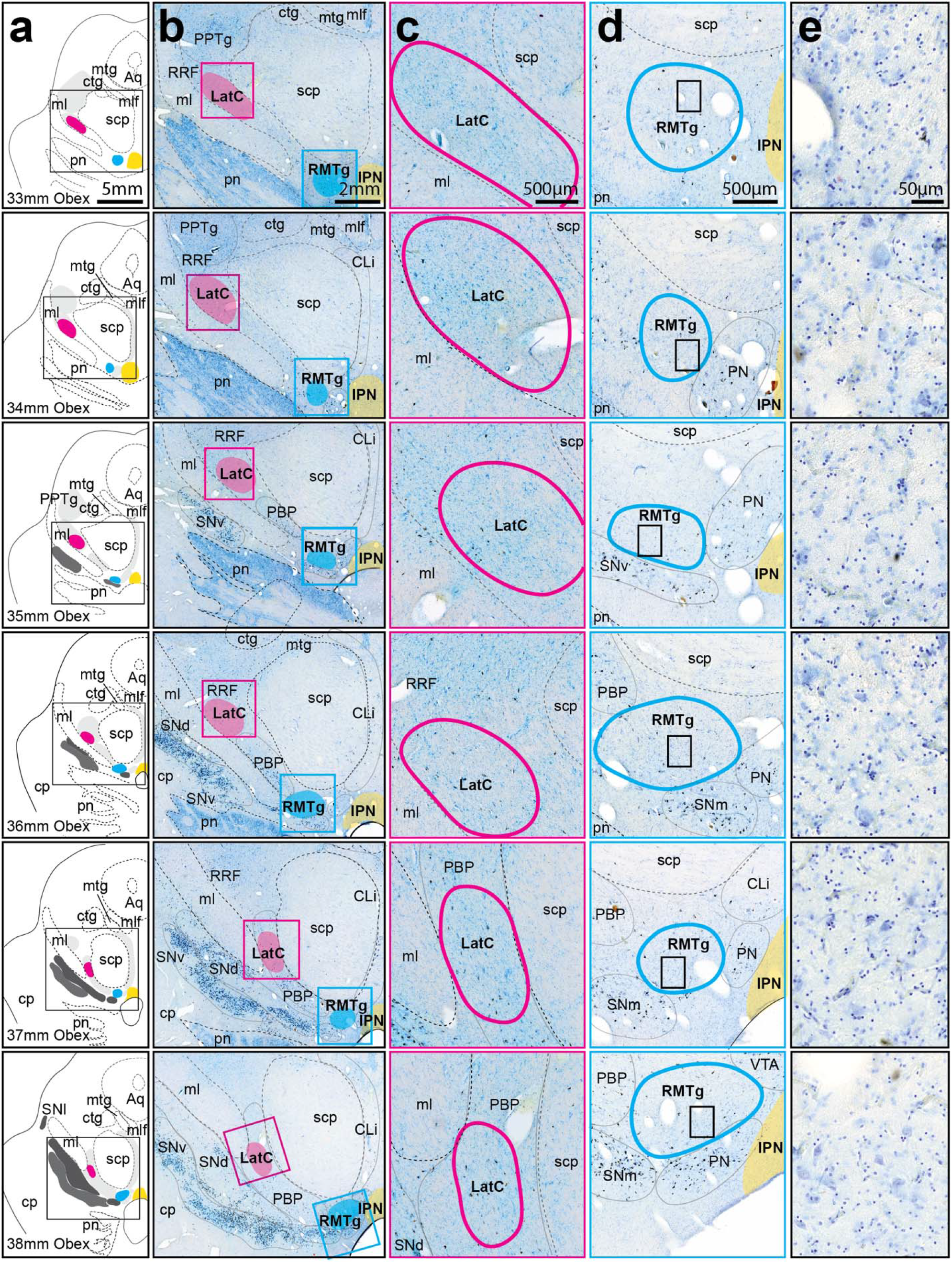
Cytoarchitectural identification of the RMTg and RRF/PBP lateral GABAergic cell cluster (LatC) in the human midbrain. a) Schematic mapping of 50μm-thick Nissl (CV) stained sections from N=4 cases with representative micrographs shown in b. Substantia nigra pars compacta (SNpc) and ventral tegmental area (VTA) subclusters, as well as retrorubral field (RRF) and other relevant nuclei (e.g. the pedunculopontine tegmental nucleus or PPTg) are annotated in grey. RRF, VTA, PPTg and several other nuclei are shown in light grey, whereas SNpc subdivisions are shaded in darker grey. The aqueduct (Aq) and fiber tracts, such as medial lemniscus (ml), superior cerebellar peduncle (scp), central tegmental tract (ctg), mammillotegmental tract (mtg) and the medial longitudinal fasciculus (mlf), are also relevant landmarks. The RRF/PBP lateral GABAergic cell cluster (LatC, magenta) was located lateral to the scp, whereas the rostromedial tegmental nucleus (RMTg, blue) was ventral to the scp and located slightly lateral to the interpeduncular nucleus (IPN, yellow). Higher magnification regional micrographs of LatC and RMTg revealed anatomical location and neighbouring structures in greater detail (c and d). Cellular view of individual RMTg neurons is presented in e. Scale bars equal 5mm in a, 2mm in b, 500μm in c and d, and 50μm in e. Other abbreviations: caudal linear nucleus (CLi); cerebral peduncle (cp); interfascicular nucleus (IF); parabrachial pigmented nucleus (PBP); paranigral nucleus (PN); pontine nuclei (pn); substantia nigra lateral part (SNl); and medial, dorsal and ventral tiers, SNm, SNd and SNv.

The cytoarchitecture of the RMTg is highlighted in more detail as this subregion has not previously been identified cytoarchitecturally in humans. In Nissl stained midbrain sections, RMTg neurons seem smaller, more round and overall denser compared to their neighboring DA cell regions (Figure 4a, d, e). Caudally the RMTg cell cluster can be identified lateral to the interpeduncular nucleus (Figure 4a, d, e, 33mm Obex), and in more rostral sections it is lateral to the paranigral nucleus (Figure 4a, d, e, 34mm, 35mm, 36mm, 37mm, 38mm Obex). Rostrally it collocates dorsal to SNpc cell clusters alongside/overlapping with other VTA DA subregions (Figure 4a, d, e, 35mm, 36mm, 37mm, 38mm Obex).

### Integrated cyto-and chemo-architectural three-dimensional (3D) model of the GABAergic cell clusters in the human RMTg and RRF/PBP

Three dimensional modelling revealed consistent overlap between the structural and chemical location of the mapped GABAergic cell clusters in both tissue types (Figure 5a). A pseudo-3D model of the RMTg and RRF/PBP LatC was constructed across cases in the rostrocaudal plane (Figure 5b). This model confirms the concentration of the RMTg to the lateral edge of the interpeduncular nucleus (Figure 5b). The larger area of the LatC in the ventromedial RRF hugs the ventrolateral edge of the superior cerebellar peduncle above and the medial lemniscus below, then travels rostomedially to the PBP and the more ventromedial aspect of the superior cerebellar peduncle (Figure 5b).

**Figure 5.**
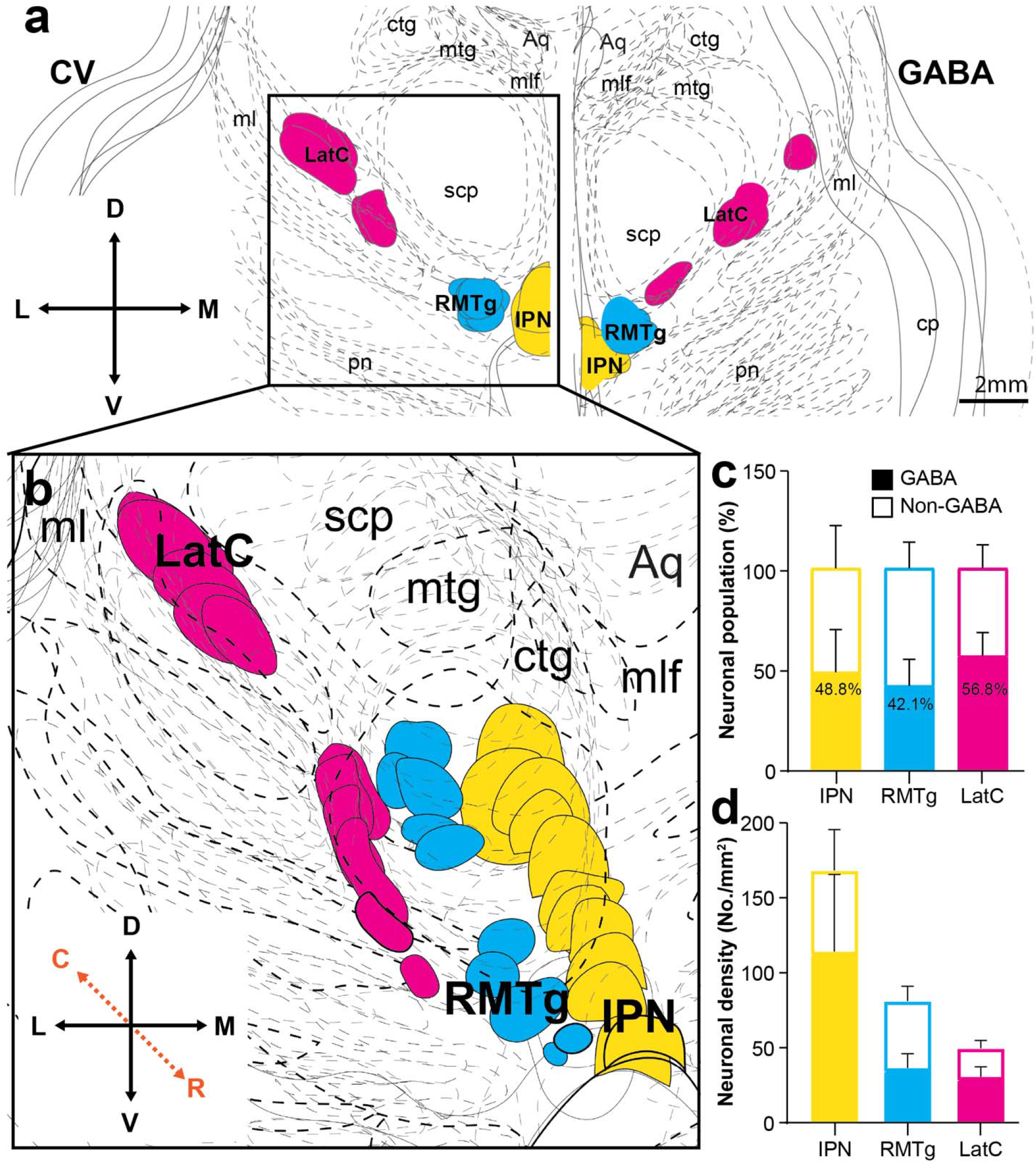
Integrated cyto- and chemo-architectural three dimensional model of the GABAergic neuronal clusters in the human midbrain DA regions. a) Schematic reconstructions of inhibitory nuclei of interest. Cross-sectional midbrain Nissl (CV) stained serial sections (left) and GABA-labeled sections (right) were collapsed and integrated across approximately 4-6mm in the rostrocaudal plane. Scale bar equals 5mm. To visualise a pseudo-3D transformation of these inhibitory midbrain nuclei, traced cytoarchitectural sections were aligned at a slight angle and the zoomed in region is shown in b. As expected, the rostromedial tegmental nucleus (RMTg, blue) closely associates with the interpeduncular nucleus (IPN), whereas the lateral cell cluster (LatC, magenta) shifts between the retrorubral field (RRF) and parabrachial pigmented nucleus (PBP) DA subregions. Within these GABAergic cell clusters, neuronal populations were further assessed in c and d in N=6 cases. Approximately half of IPN, RMTg and LatC neurons are GABAergic in c. However, the neuronal density differs with a significant main effect of anatomical region on cell density in d [F (2,28) = 3.737, * p=0.0364]. Error bars display SEM in c and d. Abbreviations: aqueduct (Aq); central tegmental tract (ctg); cerebral peduncle (cp); mammillotegmental tract (mtg); medial lemniscus (ml); medial longitudinal fasciculus (mlf); pontine nuclei (pn); superior cerebellar peduncle (scp).

The composition and density of neurons in these GABAergic cell clusters was then determined and compared. The composition of neurons was relatively uniform in the three GABAergic neuronal regions in humans with approximately 40-60% of neurons staining with the GAD67 antibody (Figure 5c). However, the density of GABAergic neurons differs between these regions with a higher density of GABAergic neurons in the interpeduncular nucleus compared to the RMTg and LatC, and a lower density of non-GABAergic neurons in the LatC [F (2,28) = 3.737, p<0.05](Figure 5d). Overall our data show that there are discrete concentrations of morphologically different GABAergic neurons in two midbrain DA regions in humans, the RMTg with smaller neurons (previously identified in non-human species), and a novel more concentrated LatC cell cluster with larger GABAergic neurons in the RRF/PBP.

## Discussion

This study identified two concentrated clusters of inhibitory neurons in the human midbrain DA regions, with one cluster equivalent to the RMTg that has been previously identified in non-human species (Figure 6a and RMTg information in Supplementary file) and the other a novel dense cluster of larger GABAergic neurons we called LatC that crosses the RRF and PBP DA subregions (Figure 6b). Morphological analyses confirmed that these two inhibitory neuronal clusters are distinct. The RMTg has not been identified in the human midbrain, and to the best of our knowledge the separate inhibitory neuronal population in the LatC has also not been identified. Approximately half of the neuronal population in these cell clusters are GABAergic with LatC having a lower density of non-GABAergic neurons compared with the RMTg. In rodents the proportion of GABAergic neurons in the RMTg is approximately 70-90% (Morello and Partanen 2015), although human brain tissue fixation and/or age may have contributed to an undercount of GABAergic neurons in our study. The average age of our longitudinally followed aged controls without neuropathological, neurological or psychiatric diseases was 85±12 years. While there is a perception that there is an age-related decline in the neuronal populations within these dopaminergic midbrain regions in humans, the most recent stereological analysis of the substantia nigra confirms previous studies showing that neuronal number (both dopaminergic and non-dopaminergic neurons) remain stable with age (Di Lorenzo Alho et al. 2016). Further, in the cognitively intact oldest-old cortical GABA levels remain stable (Britton et al. 2025), suggesting that age is unlikely to have impacted our results. We used a GAD67 antibody to label cytoplasmic GABA and longer fixation can alter GAD epitopes to reduce the number of detectable GABAergic neurons (Howat and Wilson 2014; Vincek et al. 2003). However, we used antigen retrieval methods and the proportion of non-DA to DA neurons in the substantia nigra is known to be higher in rodents compared with humans (Hardman et al. 2002), with our data consistent with this orthogonal observation. Further studies are required to confirm our findings in humans and to determine the identity of the non-GABAergic neuronal population in these midbrain nuclei.

**Figure 6.**
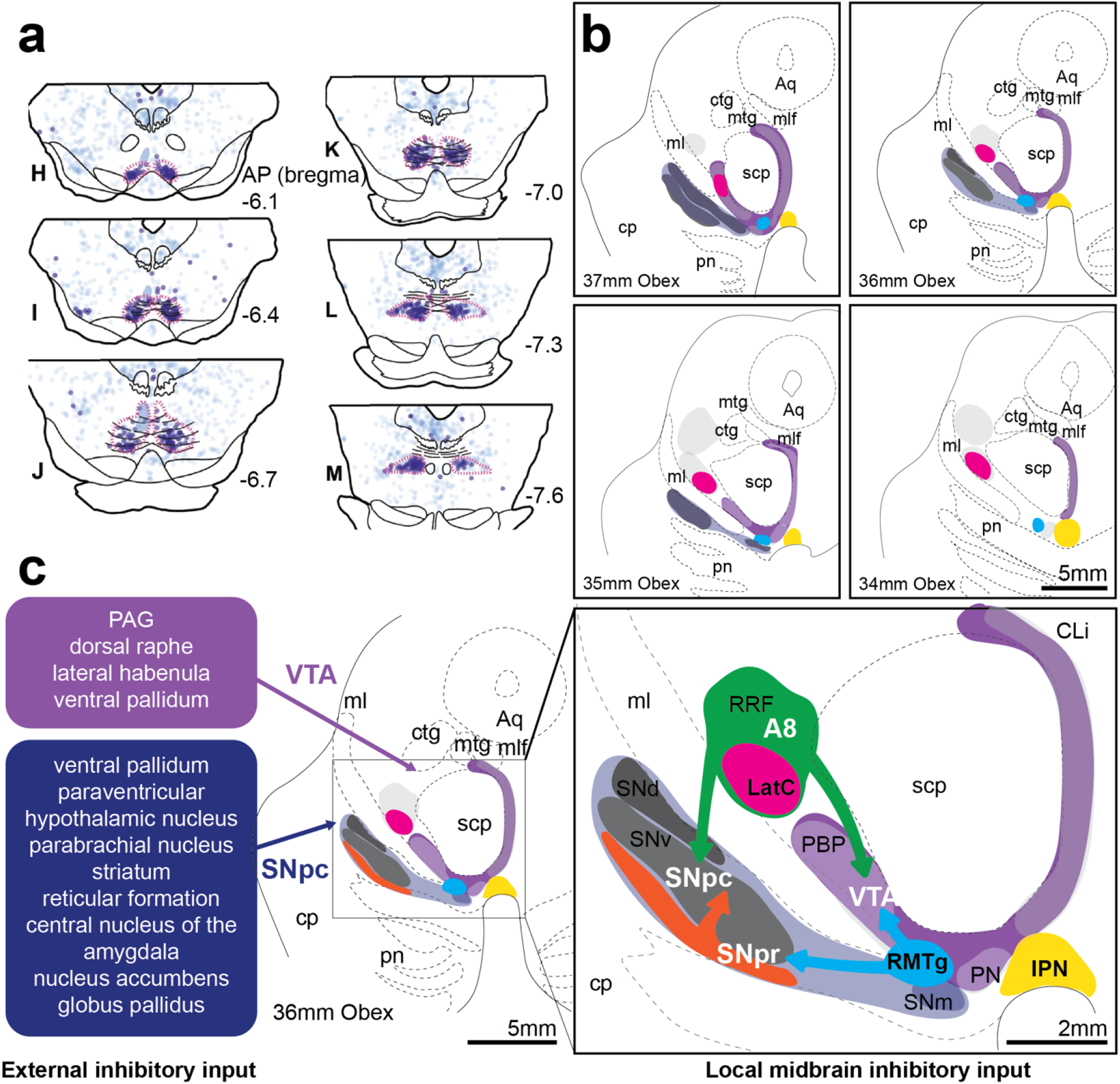
Connectivity of inhibitory nuclei to DA brain regions in humans. a) Rodent anatomical mapping of RMTg (red outlines of dark blue neuron representative circles among other tract traced neurons in light blue). Figure adapted and reproduced with permission from *Smith et al., 2019* (Smith et al. 2019). b) GABA and DA clusters in the caudal human midbrain showing the lateral cluster (LatC), RMTg and interpeduncular nucleus (IPN). c) Summarised inhibitory input from rodent and non-human primate connectivity literature. VTA receives input from periaqueductal gray, dorsal raphe, lateral habenula and ventral pallidum. SNpc also receives projections from the ventral pallidum, as well as other regions included in the blue box. These DA regions also receive inhibitory projections from the RMTg, substantia nigra pars reticulata (SNpr) and RRF. Scale bars equal 5mm in b and left side c, and 2mm in right side in c. Other abbreviations: aqueduct (Aq); caudal linear (CLi); central tegmental tract (ctg); cerebral peduncle (cp); mammillotegmental tract (mtg); medial lemniscus (ml); medial longitudinal fasciculus (mlf); parabrachial pigmented nucleus (PBP); paranigral nucleus (PN); pontine nuclei (pn); substantia nigra pars compacta (SNpc) lateral part, medial, dorsal and ventral tiers, SNl, SNm, SNd and SNv, respectively; superior cerebellar peduncle (scp).

In addition to their morphological differences, these two GABAergic nuclei have been shown to be molecularly distinct developmentally in rodents (Lahti et al. 2016; Düdükcü et al. 2024). Specifically, one study has shown that *Sox14* and *Zfmp2* transcription factors distinguish developing RMTg from RRF GABAergic neurons (Lahti et al. 2016), while another study shows that *Nkx2*.*2* identifies three developing GABAergic subtypes in the lateral non-SNpc regions (RRF not identified directly) and that *Slc32a1* identified six developing GABAergic subtypes in the SNpc and VTA regions (two GABAergic neuron subtypes and four DA+GABA neuron subtypes) (Düdükcü et al. 2024). This suggests that there are molecular differences between these midbrain GABAergic cell clusters which require further evaluation in humans.

Using anatomical landmarks, we have generated three dimensional cyto- and chemo-architectural maps of the human midbrain to show the location of these inhibitory cell clusters within the DA subregions. Mapping these midbrain inhibitory nuclei in humans will enable further understanding of their molecular signatures, function and dysfunction in disease. The recent evidence that chronic hyperactivity of DA neurons in animal models leads to DA cell dysfunction and loss (Rademacher et al. 2025) makes identifying and evaluating these midbrain GABAergic nuclei important.

### Differences between the midbrain inhibitory circuits

As identified in the introduction, midbrain DA neurons in the SNpc and VTA receive major GABA input from a variety of sources that can be divergent in function (Figure 6c). Midbrain projecting external inhibitory circuits differ functionally, molecularly and structurally. Genetically defined globus pallidus and dorsal striatum cells inhibit SNpc DA neurons but only striatal inputs induce rebound firing (Evans et al. 2020). Aldh1a1+ SNpc neurons receive inputs from molecularly distinct striatal clusters, which have opposite effects on locomotion (Dong et al. 2025), potentially due to their location on distal dendrites - striosomal GABAergic axons have a positive relationship between inhibition strength and distance away from the SNpc proximal dendrite (Evans et al. 2020). Evidence of functional differences in the input of GABA from the RMTg versus extrinsic sources to the VTA is that activation of RMTg GABA terminals induces place avoidance whereas PAG GABA terminal activation induces immobility (St. Laurent et al. 2020). Further work on determining how different inhibitory circuits influence DA neuronal function is required. Importantly, the inhibition exerted by RMTg GABAergic neurons on midbrain DA neurons is bigger compared to the one mediated for example by the substantia nigra pars reticulata GABA neurons (Morello and Partanen 2015), further warranting the study of these inhibitory midbrain nuclei in humans.

The identified RMTg and RRF/PBP LatC GABAergic nuclei must be functionally different from each other as well as external GABA inputs based on their location within different functional brain regions (Figure 6c). The GABAergic nuclei embedded in the DA subregions likely mediate different effects on midbrain DA neurons. Recent evidence in primates show that RRF and PBP GABAergic neurons receive equivalent GABAergic inputs to their DA neighbours (Fudge et al. 2024). Neural circuits with such mutual inhibitory connections are able to rapidly and flexibly switch between distinct functions allowing multiple controllable functions to be managed through the same circuits (Liu et al. 2025). The demonstration of these two distinct GABAergic nuclei that receive similar afferents to their DA neighbours and innervate broadly across all DA subregions is likely to at least double the tunable flexibility of the multiple functions regulated through DA circuits. These and other inhibitory nuclei are promising anatomical correlates for unexplored biological and disease mechanisms.

Overall, this is the first study to identify RMTg and RRF/PBP LatC GABAergic nuclei in the human midbrain. The data presented show clear concentrations of large GABAergic neurons in these human midbrain regions, higher numbers than expected for the small intrinsic interneurons also found in DA subregions. Given their major input to midbrain DA neurons it will be important to understand how their interactions impact midbrain DA motor and behavioural functions, particularly as chronic hyperactivity of DA neurons in animal models leads to DA cell dysfunction and loss (Rademacher et al. 2025).

## Acknowledgements

The authors acknowledge: Anahid Ansari Mahabadian for research assistance; Heidi Cartwright for assistance with the figures; human brain tissue sample procurement by the Sydney Brain Bank, Sydney (enabled by funding from Neuroscience Research Australia and is a special gift in memory of Jim Raftos from the Shaw family) and Flinders Medical Centre and Institute of Medical and Veterinary Sciences, Adelaide; the use of Microscopy Australia infrastructure at the Sydney Microscopy and Microanalysis facility, University of Sydney (enabled by the National Collaborative Research Infrastructure Strategy); and funding support by Aligning Science Across Parkinson’s [020505] through the Michael J. Fox Foundation for Parkinson’s Research and the National Health and Medical Research Council of Australia to GMH [Investigator Grant 2034292]. For open access, the authors have applied a CC BY public copyright license to all Author Accepted Manuscripts arising from this submission.

## Statements & Declarations

### Author contributions

All authors contributed to the study conception and design. Material preparation and data collection were performed by Anastasia Filimontseva and analysis were performed by all authors. The first draft of the manuscript was written by Anastasia Filimontseva and all authors commented on previous versions of the manuscript. All authors read and approved the final manuscript.

### Data availability

The data and analysis images, key lab materials and protocols used and generated in this study are listed in a Key Resource Table alongside their persistent identifiers at Zenodo DOI 10.5281/zenodo.17946370.

### Ethics approval

Tissue was obtained from the Sydney Brain Bank (UNSW Sydney Human Research Ethics Committee approval HC200026) and the study approved by the University of Sydney Human Research Ethics Committee (2021/845).

