## Supplementary Information for "Location and mapping of the human rostromedial tegmental nucleus and associated midbrain inhibitory nuclei regulating dopamine neurons"

### Supplementary Methods

#### Semi-automated QuPath quantification for cell typing

Morphological features of GABA and Nissl positive neurons were assessed using QuPath (ver.0.5.1, RRID:SCR_018257; https://qupath.github.io/). A semi-automated quantification method was devised to assess morphological features, cellular density and cell-type proportions in GABAergic cell clusters.

1. GABA neuronal area and circularity

GABA positive neurons in immunohistochemically processed GAD67 and TH stained sections (33.5mm – 38mm, Figure 2) were quantified with QuPath. Three separate 'pixel classifiers' were created for this analysis for each ROI - the interpeduncular nucleus (IPN), rostromedial tegmental nucleus (RMTg) and lateral cluster in the RRF (LatC). Positive neuronal staining was manually traced in ROIs in several sections. These traced images were used to 'train a pixel classifier' for each ROI. In some instances, training images were modified to increase the suitability of a pixel classifier. Once a pixel classifier showed suitable live prediction output, a size threshold was set. For GABAergic neurons this was set to a minimum 250μm$^2$ and applied across all clusters.

Our GABA antibody (GAD67, RRID:AB_627650, Cat# sc-28376) abundantly labelled GABA positive neurons, but also labelled fibers and terminals. Some cases had more intense fiber and terminal GABA labelling, which challenged the neuronal-GABA pixel classifier. As a result, the neuronal-GABA pixel classifier output was manually assessed for every case. In sections where the classifier was not sufficient, GABA-positive cells were traced manually to assess cellular morphology in these clusters. Relevant cellular morphology features were extracted for further statistical testing.

2. Nissl neuronal area and circularity

CV-stained, serial sections from relevant Obex levels (33mm – 38mm, Figure 3) were analysed in QuPath using the same methods in the three ROIs. When the pixel classifier for Nissl stained neurons with a nucleolus showed suitable live prediction output, the size threshold was set. For Nissl stained neurons this was set to a minimum of 70μm$^2$. This size threshold was consistent for all three ROIs and was selected to limit identification of non-neuronal Nissl positive nuclei. These classifiers were run on the relevant ROIs in all sections analysed. Relevant cellular morphology features were extracted for further statistical testing.

3. Neuronal cell-typing analysis

Similar pixel classifier analysis was used to investigate GABA and non-GABA neuronal populations in sections stained immunohistochemically for GAD67 and TH and then CV. After de-coverslipping and re-hydration, the sections GAD67 and TH stained sections were counterstained with 0.5% CV (HB5608 Hello Bio) to analyse the non-GABA neuronal proportion. Using the same pixel classification protocol, appropriate classifiers were created, visually validated and applied to the clusters of interest. The minimum size threshold for neuronal Nissl was set at 50μm$^2$ for all regions in all sections. After extracting the total number of non-GABA neurons, we quantified total neuronal population in these regions by adding the previously quantified GABA-positive neurons from the same sections. We then performed statistical analysis for proportion and density of GABA and non-GABA cell types.

### Supplementary Tables 1 and 2

### **Supplementary Table 1**. Case details

| **Brain Bank** | **ID** | **Age** | **Sex**# | **ABC scores*** | **Analysis** |
|---|---|---|---|---|---|
| Sydney Brain Bank, Australia | MJFF Case 3 | 85 | F | 2,0,1 | GABA neuronal morphology and density, chemoarchitectural mapping and neuronal cell-typing analysis. |
| Sydney Brain Bank, Australia | MJFF Case 5 | 88 | F | 2,1,3 | GABA neuronal morphology and density, chemoarchitectural mapping and neuronal cell-typing analysis. |
| Sydney Brain Bank, Australia | MJFF Case 11 | 91 | F | 1,2,0 | GABA neuronal morphology and density, chemoarchitectural mapping and neuronal cell-typing analysis. |
| Sydney Brain Bank, Australia | MJFF Case 12 | 93 | F | 1,1,0 | GABA neuronal morphology and density, chemoarchitectural mapping and neuronal cell-typing analysis. |
| Sydney Brain Bank, Australia | MJFF Case 7 | 97 | M | 1,1,0 | Chemoarchitectural mapping. |
| Sydney Brain Bank, Australia | MJFF Case 8 | 100 | F | 0,1,0 | Chemoarchitectural mapping. |
| Sydney Brain Bank, Australia | MJFF Case 9 | 87 | M | 2,1,0 | Chemoarchitectural mapping. |
| Sydney Brain Bank, Australia | MJFF Case 10 | 76 | F | 0,1,0 | Chemoarchitectural mapping. |
| Sydney Brain Bank, Australia | MJFF Case 4 | 93 | M | 2,0,1 | GABA neuronal morphology and density, chemoarchitectural mapping, neuronal cell-typing and supplementary figure 4 cellular cluster identification. |
| Sydney Brain Bank, Australia | MJFF Case 6 | 93 | F | 2,1,1 | GABA neuronal morphology and density, chemoarchitectural mapping and neuronal cell-typing analysis. |
| Flinders Medical Centre and Institute of Medical and Veterinary Sciences in South Australia | FMC Control 1 | 76 | F | 0,0,0 | Nissl cellular morphology and cytoarchitectural mapping. |
| Flinders Medical Centre and Institute of Medical and Veterinary Sciences in South Australia | FMC Control 2 | 58 | M | 0,0,0 | Nissl cellular morphology and cytoarchitectural mapping. |
| Flinders Medical Centre and Institute of Medical and Veterinary Sciences in South Australia | FMC Control 3 | 88 | F | 0,1,0 | Nissl cellular morphology and cytoarchitectural mapping. |
| Flinders Medical Centre and Institute of Medical and Veterinary Sciences in South Australia | FMC Control 4 | 64 | M | 0,0,0 | Nissl cellular morphology and cytoarchitectural mapping. |

#M=male, F=female; *(Hyman et al. 2012)

**Supplementary Table 2**. Rostromedial tegmental nucleus (RMTg) identification in animal studies to identify relevant landmarks.

| **Species** | **Rostral landmarks** | **Intermediate landmarks** | **Caudal landmarks** | **Method** | **Selective RMTg labeling** | **Reference** |
|---|---|---|---|---|---|---|
| Rat | | CLi, IPN, xscp | | Retrograde injections in VTA and lateral habenula to label RMTg neurons | Yes | (Schoukroun et al. 2024) |
| Rat | ml, IPN | IPN, scp | xscp | FosB and GABA double labeling after chronic cocaine administration | Yes | (Perrotti et al. 2005) |
| Rat | NA | NA | NA | VTA-injected retrogradely labeled RMTg neurons co-expressing GAD67; and retrogradely labeled Fos positive neurons | Yes | (Jhou et al. 2009a) |
| Rat | Posterior VTA | NA | Xscp, cp, ml, IPN, tth | FosB immunoreactivitiy with psychostimulant injections | Yes | (Kaufling et al. 2010) |
| Rat | Posterior VTA and PN, ml, IPN, CLi, RRF, RN, cp | xscp, CLi, IPN, ml, tth, RRF, cp | xscp, CLi, IPN, Pn, cp, ml, tth, RRF | Double immunostaining for FosB and GABA | Yes | (Kaufling et al. 2009) |
| Rat | CLi, cp, RRF, IPN | NA | NA | Klüver-Barrera stain | No | (Petzel et al. 2017) |
| Mouse | NA | NA | NA | Anterograde injection into lateral habenula and GAD67 immunofluorescence | No | (Lammel et al. 2012) |
| Mouse | NA | NA | NA | Sox14 and FoxP1 *in-situ* hybridization and immunohistochemical labeling in embryonic mice | Yes | (Lahti et al. 2016) |
| Mouse | IPN, VTA, RLi | MnR, CLi | MnR | Viral injections encoding muscarinic Gq protein-coupled receptors (hM3Dq) injected into RMTg | Yes | (Zhao et al. 2022) |
| Non-human primates | | xscp, ml, mlf, IPN, NRTP | | Electrophysiological recordings of orthodromic and antidromic stimulations of lateral habenula and SNpc. Recorded RMTg neurons were histologically reconstructed. RMTg boundaries were delineated with retrograde tracing. | | (Hong et al. 2011) |

CLi=caudal linear nucleus, cp=cerebral peduncle, IPN=interpeduncular nucleus, ml=medial lemniscus, mlf=medial ongitudinal fasciulus, MnR=median raphe nucleus, NRTP=nucleus reticularis tegmenti pontis , Pn=pontine nuclei, PN=paranigral nucleus, RLi=rostral linear nucleus, RN=red nucleus, RRF=retrorubral fields, scp=superior cerebellar peduncle, SNpc=substantia nigra pars compacta, tth=trigeminothalamic tract, VTA=ventral tegmental area, xscp=decussation of the scp

### Supplementary Figures 1-3

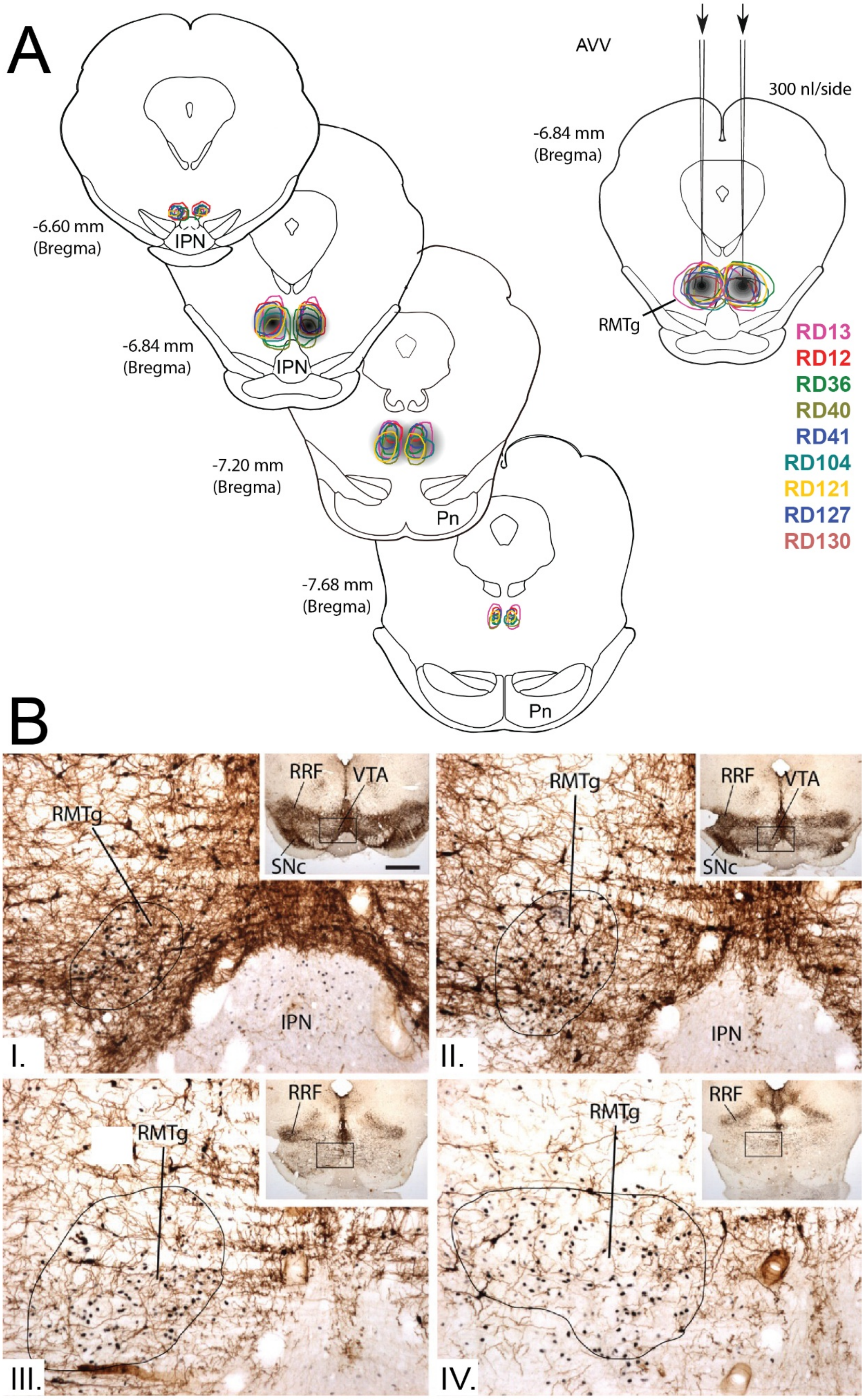

**Supplementary Figure 1.** ***Identification and mapping of rostromedial tegmental nucleus (RMTg) in rodent using stereotaxic surgery and immunohistochemistry.*** **A)** Adeno-associated virus (AAV) vectors expressing hM3Dq receptors targeted RMTg at different rostro-caudal levels in nine rats. Figure adapted with permission from *Yang et al., 2018* (Yang et al. 2018). **B)** RMTg neurons also express Fos protein (black reaction product) after methamphetamine injection. Fos-labeled neurons are embedded in tyrosine hydroylase (TH) immunoreactive processes and cell bodies (brown) in more rostral sections, I) and II). There is less TH immunoreactivity in caudal sections, III) and IV). The RMTg outline is larger in more caudal sections. Figure adapted with permission from *Jhou et al., 2009* (Jhou et al. 2009b). Abbreviations: aqueduct (Aq); pontine nuclei (Pn); retrorubral field (RRF); interpeduncular nucleus (IPN); substantia nigra pars compacta (SNc); ventral tegmental area (VTA).

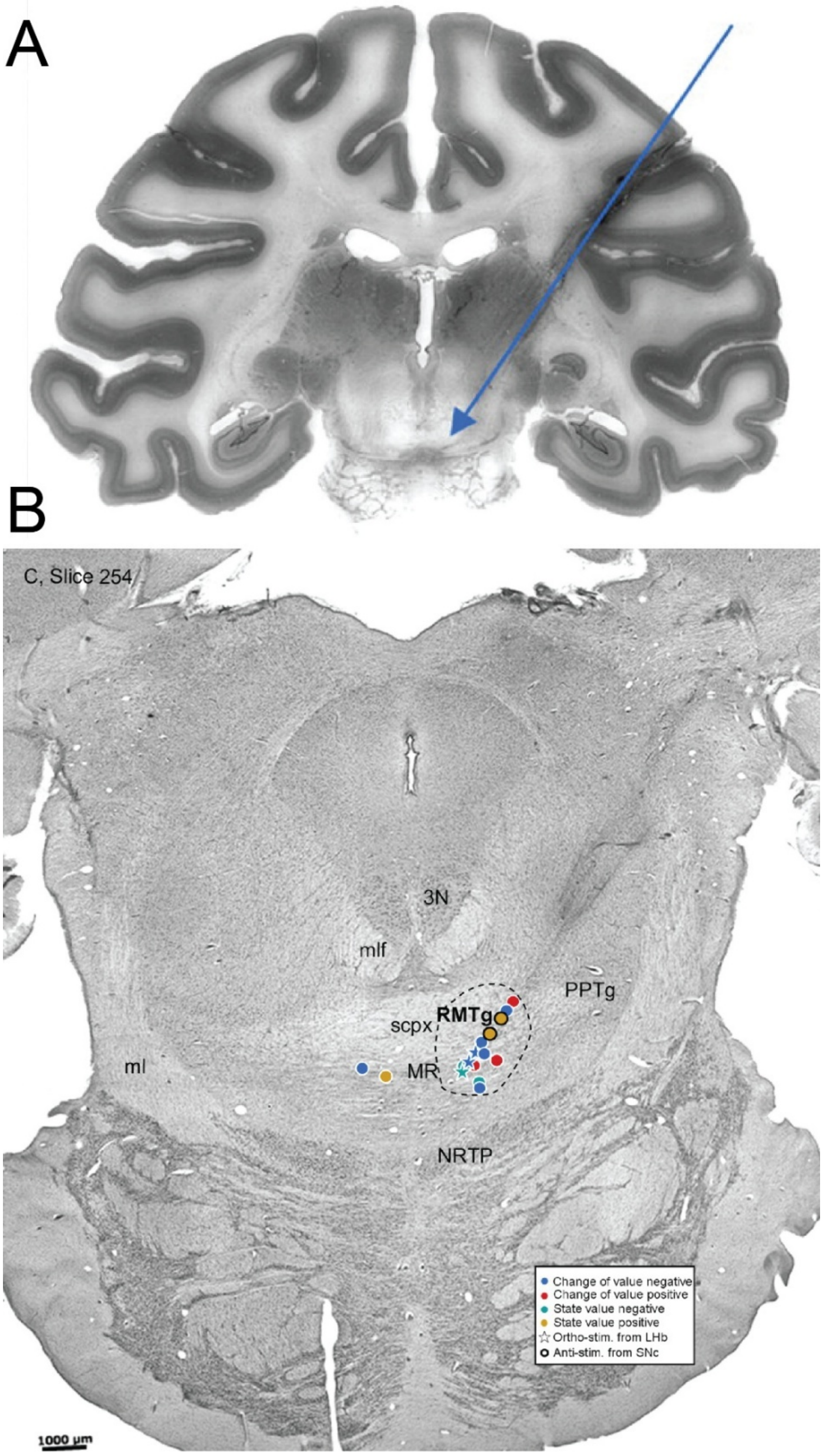


**Supplementary Figure 2.** ***Identification of rostromedial tegmental nucleus (RMTg) in rhesus monkeys.*** **A)** Electrode tract schematic to identify RMTg neurons during a reward-related visual task. **B)** Identity of recorded neurons was confirmed as RMTg using orthodromic and antidromic stimulations from lateral habenula and substantia nigra pars compacta (SNpc), respectively. Estimated locations of recorded neurons are histologically mapped. Figure adapted with permission from *Hong et al., 2011* (Hong et al. 2011). Abbreviations: oculomotor nucleus (3N); medial lemniscus (ml); medial longitudinal fascilicus (mlf); median raphe (MR); pedunculopontine tegmental nucleus (PPTg); superior cerebellar peduncle decussation (scpx); nucleus reticularis tegmenti pontis (NRTP).

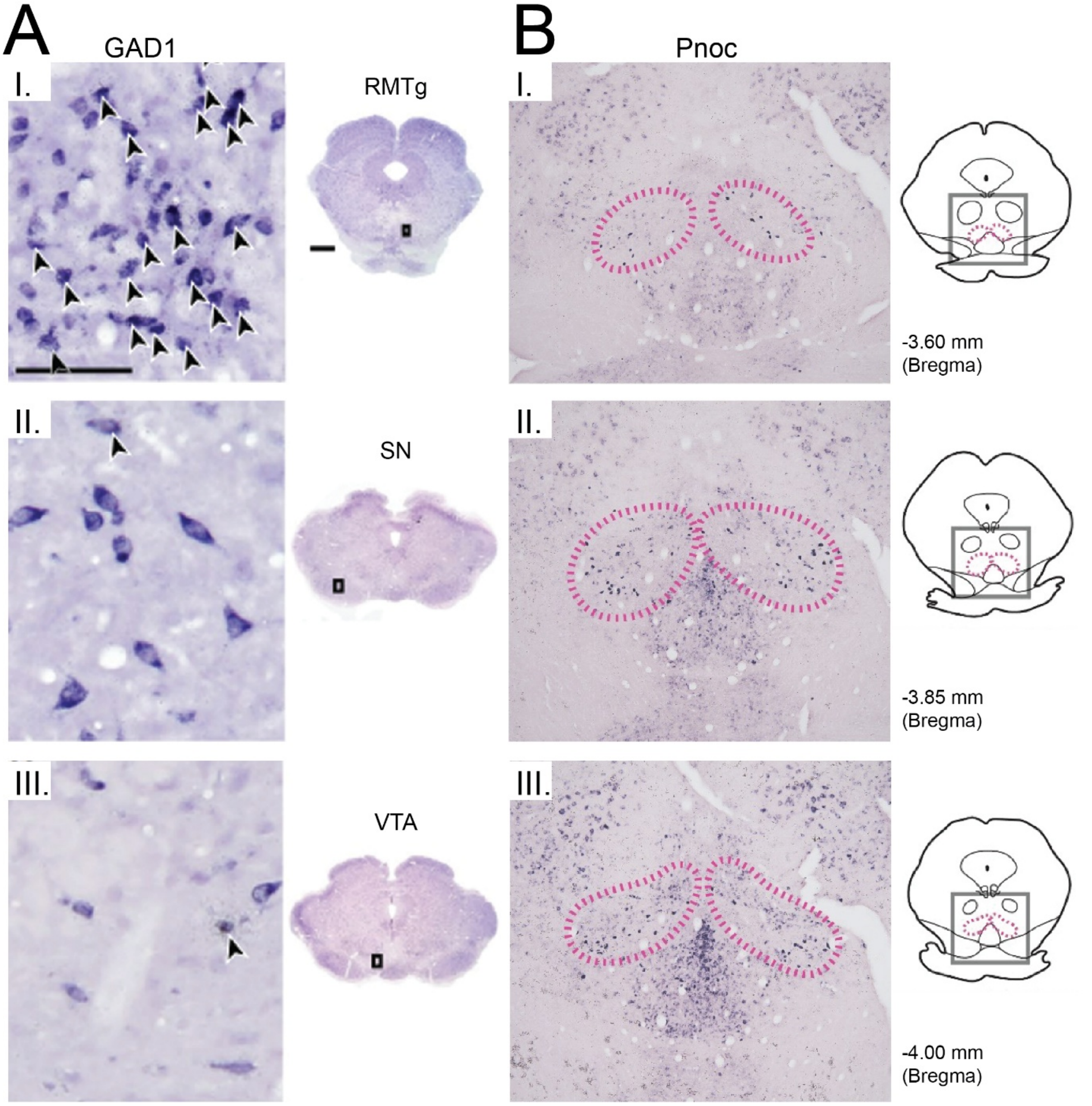


**Supplementary Figure 3.** ***Comparison of rostromedial tegmental nucleus (RMTg) neurons to neighbouring structures using in-situ hybridization.*** **A)** Inhibitory neurons in RMTg, substantia nigra (SN) and ventral tegmental area (VTA) are labelled with *GAD1 in-situ* hybridization. RMTg neurons are smaller and more dense in I), compared to SN and VTA neurons in II) and III). **B)** RMTg neurons also appear more dense and larger than neighbouring interpeduncular nucleus (IPN) when labeled with *in-situ* hybridization for *Pnoc* mRNA. Figure adapted permission from *Smith et al., 2019* (Smith et al. 2019)
